# Structural conservation of MALAT1 long non-coding RNA in cells and in evolution

**DOI:** 10.1101/2022.07.29.502018

**Authors:** Anais Monroy-Eklund, Colin Taylor, Chase Weidmann, Christina Burch, Alain Laederach

**Affiliations:** Department of Biology, University of North Carolina at Chapel Hill, Chapel Hill, NC, 27599; Department of Biological Chemistry, University of Michigan Medical School, Center for RNA Biomedicine, Rogel Cancer Center, Ann Arbor Michigan, MI, 48103

## Abstract

Although not canonically polyadenylated, the long non-coding RNA MALAT1 (Metastasis Associated Lung Adenocarcinoma Transcript 1) is stabilized by a highly conserved 159 nucleotide triple helix structure on its 3’ end. The entire MALAT1 transcript is over 8,000 nucleotides long in humans and is considered one of the most conserved lncRNAs, at both the sequence and structure levels. The strongest structural conservation signal (as measured by co-variation of base-pairs) is in the triple helix structure. Primary sequence analysis of co-variation alone cannot confirm the degree of structural conservation of the entire full-length transcript. Furthermore, RNA structure is often context dependent; RNA binding proteins that are differentially expressed in different cell types may alter structure. We investigate here the in cell and cell free structures of the full-length human and green monkey (*Chlorocebus sabaeus*) MALAT1 transcripts in multiple tissue-derived cell lines using SHAPE chemical probing. Our data reveals surprising levels of uniform structural conservation in different cell lines, in cells and cell free, and even between species, despite significant differences in primary sequence. The uniformity of the structural conservation across the entire transcript suggests that, despite seeing co-variation signals only in the three-helix junction of the lncRNA, the rest of the transcript’s structure is remarkably conserved at least in primates and across multiple cell types and conditions.

## Introduction

MALAT1 (Metastasis Associated Lung Adenocarcinoma Transcript 1) is an ∼8kb long non-coding RNA (lncRNA) (Ji et al. 2003). It is one of the few highly expressed and highly conserved long non-coding RNAs with multiple functions in both tumors and normal cells (Hutchinson et al. 2007; Ulitsky and Bartel 2013; Arun et al. 2020). MALAT1 is conserved both at the sequence and RNA secondary structure level (Zhang et al. 2017; Tavares et al. 2019; Rivas 2021). MALAT1 has a known and conserved triple helical tertiary structure on its three-prime end (Wilusz et al. 2008; Brown et al. 2012; Wilusz et al. 2012; Brown et al. 2014; Zhang et al. 2017). Additionally, co-variation analysis has supported other specific secondary structures near the triple helical region (Zhang et al. 2017; Rivas 2021). As a result, a careful in cell RNA structural characterization of full length MALAT1 has the potential to reveal previously unknown RNA structures in the rest of the transcript. In addition, it can reveal the role of the cellular environment in modulating the underlying structure.

The size of MALAT1 (∼8kb) makes it quite similar in length to many human messenger RNAs (Waldern et al. 2021). It is transcribed by RNA Polymerase II, but, unlike mRNAs in the cell, MALAT1 is largely retained in the nucleus and is not actively translated by the Ribosome (Hutchinson et al. 2007; Tripathi et al. 2010; Djebali et al. 2012; Ingolia et al. 2014). MALAT1 undergoes noncanonical processing; RNase P cleaves the MALAT1 three prime end to produce a Trna like small molecule called mascRNA and the mature MALAT1 transcript. MALAT1 mature sequence is not polyadenylated but rather has a genetically encoded A-rich end that is critical in the formation of the triple helix (Brown et al. 2012; Wilusz et al. 2012; Brown et al. 2014). The cleaved MALAT1 three prime end folds back onto itself, creating the triple-helical architecture resistant to exonuclease-mediated decay (Brown et al. 2012; Wilusz et al. 2012; Brown et al. 2014). Thus, mature MALAT1 lncRNA is surprisingly stable and is highly expressed in broad range of cell types (Hutchinson et al. 2007; Zhang et al. 2012).

MALAT1 has many proposed functions in the cell ranging from pre-mRNA regulation of splicing to transcriptional regulation, including interacting with transcriptional repressive complex PRC2, interacting with transcription factors, and acting as a competing endogenous RNA (ceRNA) (recently reviewed by (Arun et al. 2020)). It is generally over expressed in tumors, however, a Malat1 deletion resulted in a reduction in lung metastases in one study (Gutschner et al. 2013), while another study with a Malat1 knockout promoted metastasis (Kim et al. 2018). Additionally, murine knock-out studies of Malat1 have shown that it is nonessential for life (Nakagawa et al. 2012; Zhang et al. 2012). Although MALAT1 has been extensively studied, it’s endogenous and oncogenic function has yet to be fully elucidated.

Chemical probing techniques allow us to probe RNA structure with single nucleotide resolution in a wide variety of conditions (Deigan et al. 2009; Hajdin et al. 2013; Woods and Laederach 2017; Lackey et al. 2018; Smola and Weeks 2018; Spasic et al. 2018). Furthermore, with the throughput of next generation sequencing, it is possible to obtain high-resolution quantitative data even on large transcripts (Kutchko et al. 2018; Lackey et al. 2018; Mustoe et al. 2018; Smola and Weeks 2018). It is important to distinguish between transcriptome-wide studies, where low to medium coverage provides semi-quantitative results on all transcripts, and high precision targeted amplification strategies that offer quantitative chemical probing reactivities across entire transcripts (Weeks 2015; Smola et al. 2016; Busan and Weeks 2017; Woods et al. 2017). In this study, we obtain high-resolution quantitative SHAPE-MaP (Selective 2’ Hydroxyl Acylation by Primer Extension with Mutational Profiling) on the MALAT1 lncRNA to investigate surprisingly subtle changes in RNA structure across three cell lines in two species (Siegfried et al. 2014; Mustoe et al. 2018; Smola and Weeks 2018).

Using these data, we analyze how the cellular environment (in cell vs. cell free) affect MALAT1 lncRNA structure. We also investigate how evolutionary substitution in primates affect these structures. Our findings rely on the very high level of quantitative reproducibility of targeted SHAPE-MaP data to reveal a surprisingly robust set of structures in the MALAT1 lncRNA throughout the entire transcript. We investigate these structures to determine how well they persist both in cell and cell free and with respect to evolutionary drift. Together, our work demonstrates a surprising consistency in RNA structure in the MALAT1 lncRNA, even in regions that are not known to be essential for function. Our study therefore suggests that specific structures in very long non-coding RNAs like MALAT1 may be unaffected by changes in sequence between humans and African green monkeys.

## Results

### In cell primary structure of MALAT1 transcript

We begin our structural investigation of the MALAT1 lncRNA locus in the human genome by reporting its annotated genome coordinates and overall locus conservation in Figure 1A. As is currently reported in GRCh38 (GenBank Accession GCA_000001405.28), MALAT1 is located on chromosome 11. To display MALAT1’s conservation, we are using PhyloP data generated from comparing the genomes of 30 mammals in this case, which can show both well conserved (as indicated by positive PhyloP scores) and accelerating regions (negative scores) (Pollard et al. 2010). MALAT1 has three annotated transcript isoforms (NR_002819.4, NR_144567.1, and NR_144568.1). The full-length isoform (NR 002819.4) is unspliced, while NR_144567.1 is composed of two exons, and NR_144568.1 is composed of 3 exons. We performed RNA-seq experiments on Hek293 (ATCC CRL 1573) and A549 (ATCC CCL 185) cell lines to identify the regions of the MALAT1 transcript with the highest expression and which are readily reverse transcribed during library preparation (Supplementary Figure 1A and Figure 1B). We observed uniform coverage for NR_144567.1, however only a small proportion of reads mapped to regions corresponding to the first exon. Thus, we concluded NR_144567.1 exon 2 is the dominantly expressed isoform in the cell lines we will investigate here. Our observation of expression <100 nucleotides downstream from NR_144567.1 exon 2, is consistent with what has been previously reported (Wilusz et al. 2008; Gutschner et al. 2013). We also observed no coverage on the 3’ end of the transcript in the triple-helix and mascRNA regions (Wilusz et al. 2008; Brown et al. 2012; Wilusz et al. 2012; Brown et al. 2014). This is likely due to the high level of structure in these regions and the inability of the reverse transcriptase to transcribe these regions (Gutschner et al. 2013; Sun et al. 2017; Xiping et al. 2018). We conclude that the primary structure of the lncRNA MALAT1 is the same in the systems we will study in this manuscript.

**Figure 1:**
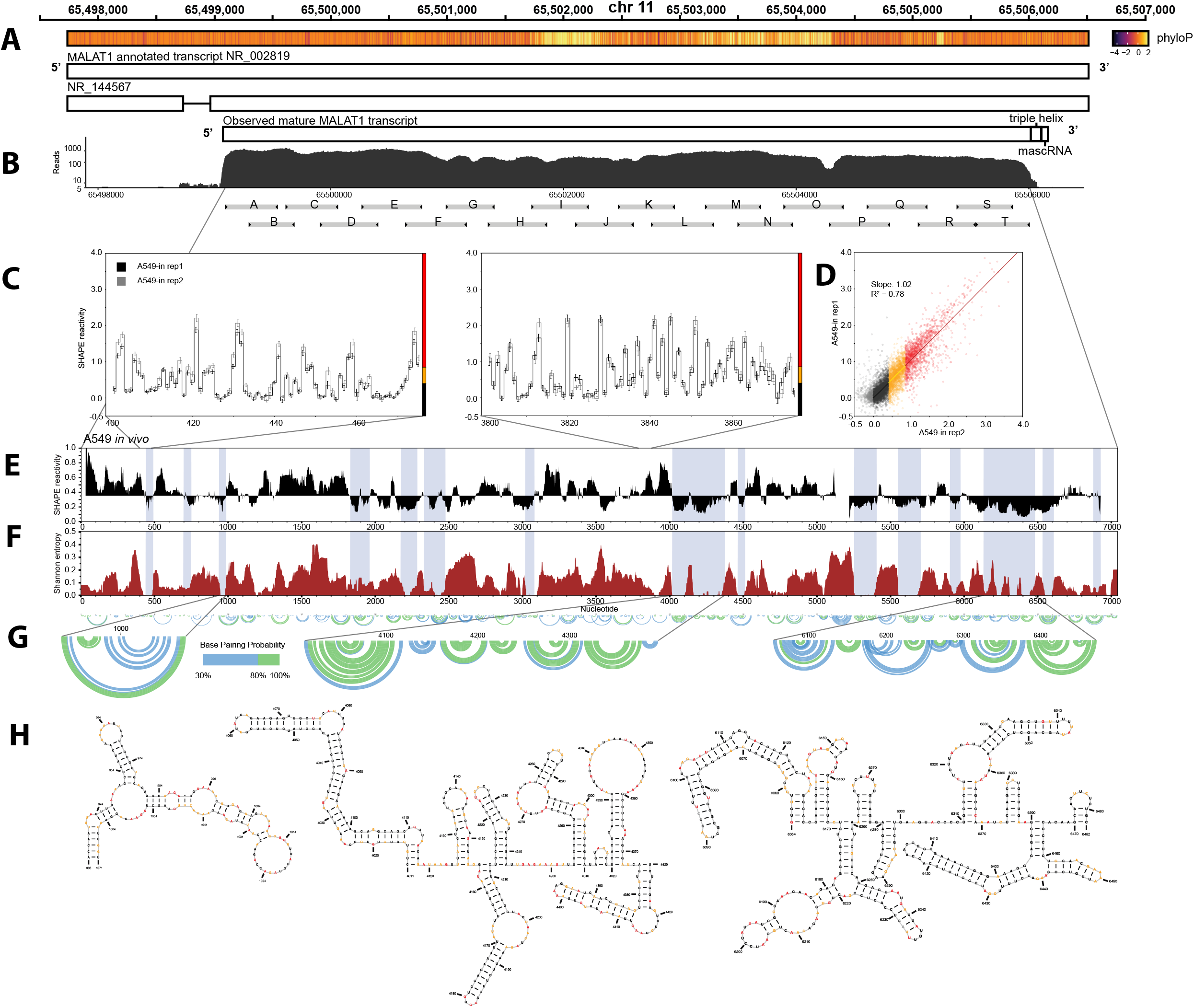
Human MALAT1 in cell structure, SHAPE-MaP reproducibility and structured regions. A) MALAT1 genomic locus, phlyoP conservation and structure for single exon NR_002819 and alternatively spliced NR_1445671 transcripts (GRCh38). B) Observed MALAT1 transcript in Hek293 cells by RNA-sequencing shows only exon 2 of NR_1445671 is expressed, with a uniform expression profile. The triple helix and mascRNA are however not detected by RNA-seq, likely due to their very high degree of structure. We designed 20 tiled and overlapping primers for amplicon amplification of SHAPE-MaP probing read through reverse transcription (labeled A-T) to cover the expressed area. C) SHAPE-MaP data is highly reproducible in cell across replicates. Replicate 1 is plotted in black, replicate 2 in gray and standard error as error bars. D) Scatter plot showing linear relationship between in cell replicates and high level of quantitative correlation (R^2^ = 0.72). Coloring is based on SHAPE reactivity convention, with low shape black, medium yellow and high red. E) Median windowed SHAPE reactivity for entire MALAT1 lncRNA used to identify regions of low SHAPE which fold to unique, well-defined structures, also shows almost complete coverage for entire lncRNA. F) Shannon Entropy of the partition function computed using SHAPE as a pseudo-free energy term in nearest neighbor modeling of RNA structure (Deigan et al. 2009). Regions of low median SHAPE and low Shannon entropy adopt well defined structures as can be seen in G) base-pairing probability arc plots for entire MALAT1 RNA structure folded using SHAPE in cell data. Select regions of low SHAPE and low Shannon entropy are enlarged and the corresponding minimum free energy structures are shown in H) The minimum free energy structure nucleotides are colored according to the low, medium, and high reactivity definitions described above, and we see that most paired nucleotides have low reactivity.

### In cell secondary structure probing and modeling

We designed a series of 20 overlapping primer pairs for selective amplicon amplification of the expressed region of the MALAT1 transcript using Selective 2’ Hydroxyl Acylation by Primer Extension with Mutational Profiling (SHAPE-MaP) (Siegfried et al. 2014; Smola et al. 2015; Lackey et al. 2018; Smola and Weeks 2018). These are labeled A-T in Figure 1B, and illustrate our overlapping strategy, with each primer pair amplifying 500-600 nucleotides of the transcript, which corresponds to the maximum read length of an Illumina sequencer. We used the SHAPE reagent 5-nitroisatoic anhydride (5NIA) to probe the cellular RNA before Trizol extraction and refolding, as outlined in the material and methods (Kutchko et al. 2015; Smola et al. 2015; Smola et al. 2016; Corley et al. 2017; Kutchko and Laederach 2017; Busan and Weeks 2018). In Figure 1C we show representative regions of our SHAPE data for two independent replicates of our in cell data collected in A549 cells. As is summarized in the scatter plot in Figure 1D, these two replicates are highly correlated and the data are quantitatively reproducible (R^2^ =0.78, slope 1.02) for low (black), medium (yellow) and high (red) SHAPE reactivities. This high reproducibility of our data is important as the subsequent analyses in this manuscript will include quantitative comparisons of these data across different conditions.

Figure 1E illustrates the median windowed SHAPE across the entire MALAT1 transcript for the combined average of both replicates, which will be used for comparisons going forward (Rice et al. 2014). It also illustrates that our primer amplicon strategy was successful at capturing high quality SHAPE data for most nucleotides spanning the MALAT1 transcript in cells. From these data we also compute the Shannon Entropy (Figure 1F) and identify regions of low SHAPE, low Shannon entropy indicated as vertical gray shading (Kennedy et al. 2008; Hamada et al. 2009; Busan and Weeks 2017). These regions are more likely to fold into a single, well defined RNA structures and be amenable to minimum free energy structural analysis (Kutchko et al. 2018; Lackey et al. 2018; Dadonaite et al. 2019). In Figure 1G we show SHAPE derived base-pairing probability arc plots for the entire transcript. The arcs observed on the SHAPE derived models of base-pairing probability indicate high support for specific base-pairs (Deigan et al. 2009; Hajdin et al. 2013; Spasic et al. 2018) and in Figure 1H we show the minimum free energy models with SHAPE data derived from the experimental reactivities (Mathews 2006; Tyagi and Mathews 2007; Deigan et al. 2009) for three representative low SHAPE, low Shannon entropy regions. Together these results show that our tiling amplicon strategy produces quantitatively reproducible in cell structure profiling data for the observed region of the MALAT1 transcript and suggests we can use this approach to further investigate the effects of cellular environment and ultimately evolutionary substitutions on the MALAT1 structure.

### Cell free structural probing MALAT1 lncRNA

In Figure 2A we focus on exon2 of the MALAT1 transcript (NR_144567.1) which we observed in our RNA-seq experiments. In Figure 2B we show that cell free SHAPE data is highly reproducible across replicates (R^2^ 0.88, slope 1.01). Here again, we used 5NIA to probe the cellular RNA, but perform the modification after Trizol extraction and refolding as outlined in the materials and methods (Kutchko et al. 2015; Weeks 2015; Corley et al. 2017; Smola and Weeks 2018). We compare cell free data replicates obtained from RNA extracted from Hek293 and A549 cells. These results show that our cell free refolding protocol is effective regardless of which cell type the RNA is extracted from, and that the MALAT1 lncRNA in the absence of proteins folds to a very similar (identical within experimental error) regardless of cell of origin. From these data we can compute the averaged median SHAPE (Figure 2C) and Shannon entropy (Figure 2D). Most of the low SHAPE, low Shannon entropy regions identified in cell overlap with those identified here cell free.

**Figure 2:**
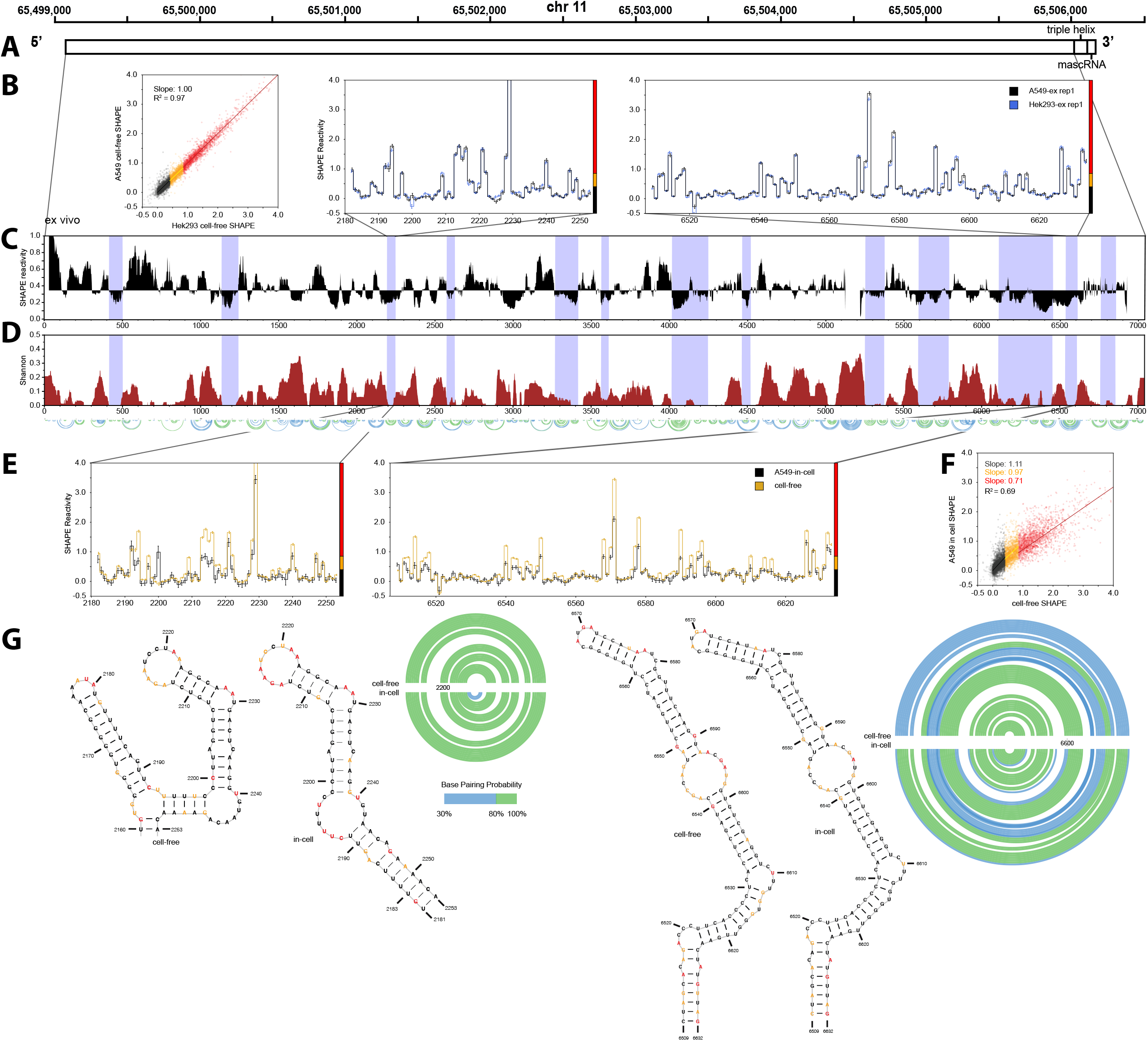
in vs. cell free MALAT1 RNA structure reveals global changes in reactivity with minimum effect on highly structured regions. A) Schematic of expressed MALAT1 transcript in human (GRCh38) genome coordinates we observed in both A549 and Hek293 cells. B) cell free SHAPE-MaP data is highly reproducible across replicates and independent of cell line origin. C) Median windowed (50 nt) SHAPE reactivity for cell free data combined with D) SHAPE derived Shannon Entropy identifies similar regions of high structure in both cell free and in cell SHAPE data sets. E) Correlation analysis of SHAPE reactivities colored by low (black), medium (yellow) and high (red) reactivity shows a clear decrease in slope for highly reactive nucleotides, despite an overall correlation comparable to in cell replicates. This is clearly visible in comparison plots of reactivities in F) where we see that the in cell data is significantly lower mostly for the highly reactive nucleotides. However, the overall pattern, which indicates structure is very similar. G) Indeed, when we model base pairing probabilities (arc diagrams) and corresponding minimum free-energy structures of low SHAPE, low Shannon entropy regions in the MALAT1 mRNA we obtain very similar structures both in and cell free. This suggests that the cellular environment is not significantly altering the structure of the MALAT1 mRNA and that the cell free observed structures persist in the cell.

When we compare the SHAPE profiles for representative regions in cells (black) with cell free (yellow) in Figure 2E we observe that the SHAPE pattern is overall the same. The nucleotides with the largest differences seem to be high SHAPE regions (colored as red, SHAPE>0.8 in the side bar), with the cell free reactivity in most cases higher than that of in cell. This is further evident in the scatter plot illustrated in Figure 2F, where the slope of the highly reactive nucleotides is lower than that of the medium (yellow) and low (black) nucleotides. As a reminder, replicates have a slope of 1.00 in Figure 2B. The quality of our data therefore allows us to observe these subtle but important global differences in SHAPE reactivity.

When we incorporate the cell free data in secondary structure models of low SHAPE, low Shannon entropy regions (shown in Figure 4G) we observe similar base-pairing probabilities and even minimum free energy structures. These results contradict some other structural probing studies of cellular RNAs (Rouskin et al. 2014; Tomezsko et al. 2020), which reported very large differences for data collected in cell and cell free conditions. It is important to note that we have collected data with high levels of reproducibility, as is evidenced by our replicate analysis in Figure 1. This requires very deep sequencing of SHAPE-MaP libraries and particular care when performing probing experiments. Interestingly, applying this approach we observe strong agreement between cell free and in cell conditions in our experiments. Nonetheless, we do observe a systematic difference, where highly reactive nucleotides tend to be less reactive in cells which is consistent with a global shielding of the chemical probe by the cellular environment. When we model the low-SHAPE low-Shannon entropy regions incorporating cell free and in cell we obtain similar structures (Figure 2G). It is the pattern of SHAPE reactivity and not its’ absolute value that directs structure prediction which explains this observation (Deigan et al. 2009; Hajdin et al. 2013; Spasic et al. 2018).

### Effect of different cellular environments on MALAT1 lncRNA structure

So far, we have established that our amplicon tiling strategy produces quantitatively reproducible data both in cells and cell free. This is essential as even when we compare in cell to cell free conditions we obtain remarkably similar SHAPE profiles. Because the underlying protein free structure of MALAT1 is unchanged regardless of cell source (Figure 2B), and because in cell probing yields very similar reactivity profiles to probing of cell free RNA (Figure 2E), we asked whether we might observe differences in lncRNA structure if we compare SHAPE data collected from in cell probing of two different cell lines. We therefore collected replicates of in cell SHAPE data in Hek293 cells, which express the same transcript isoform observed in A549 cells (Figure 3A, Supplementary Figure 1B). In these cell lines we also obtain reproducible data across two replicates (Figure 3B), from which we obtain median SHAPE profiles (Figure 3C), and identify low SHAPE, low Shannon regions that mostly overlap with those previously identified (Figure 3D). Indeed, when we compare the Hek293 averaged replicate data to A549 averaged replicate data (Figure 3E) we observe a nearly identical pattern and a high correlation (R^2^ = 0.79). We do not observe any systematic differences between low, medium, and high SHAPE slopes, but the overall slope (Figure 3E) is 0.78. This likely indicates that the 5NIA SHAPE reagent more easily permeates A549 cells, with a slightly lower overall reactivity in Hek293 cells. Nonetheless, the in cell base-pairing probabilities and corresponding minimum free energy models informed by these data are nearly identical (Figure 3F) for low SHAPE, low Shannon entropy regions. Thus, although the specific cell line affects overall SHAPE reactivity (likely due to differences in overall cell permeability of the SHAPE reagent), the structure of MALAT1 is surprisingly consistent across different cellular conditions.

**Figure 3:**
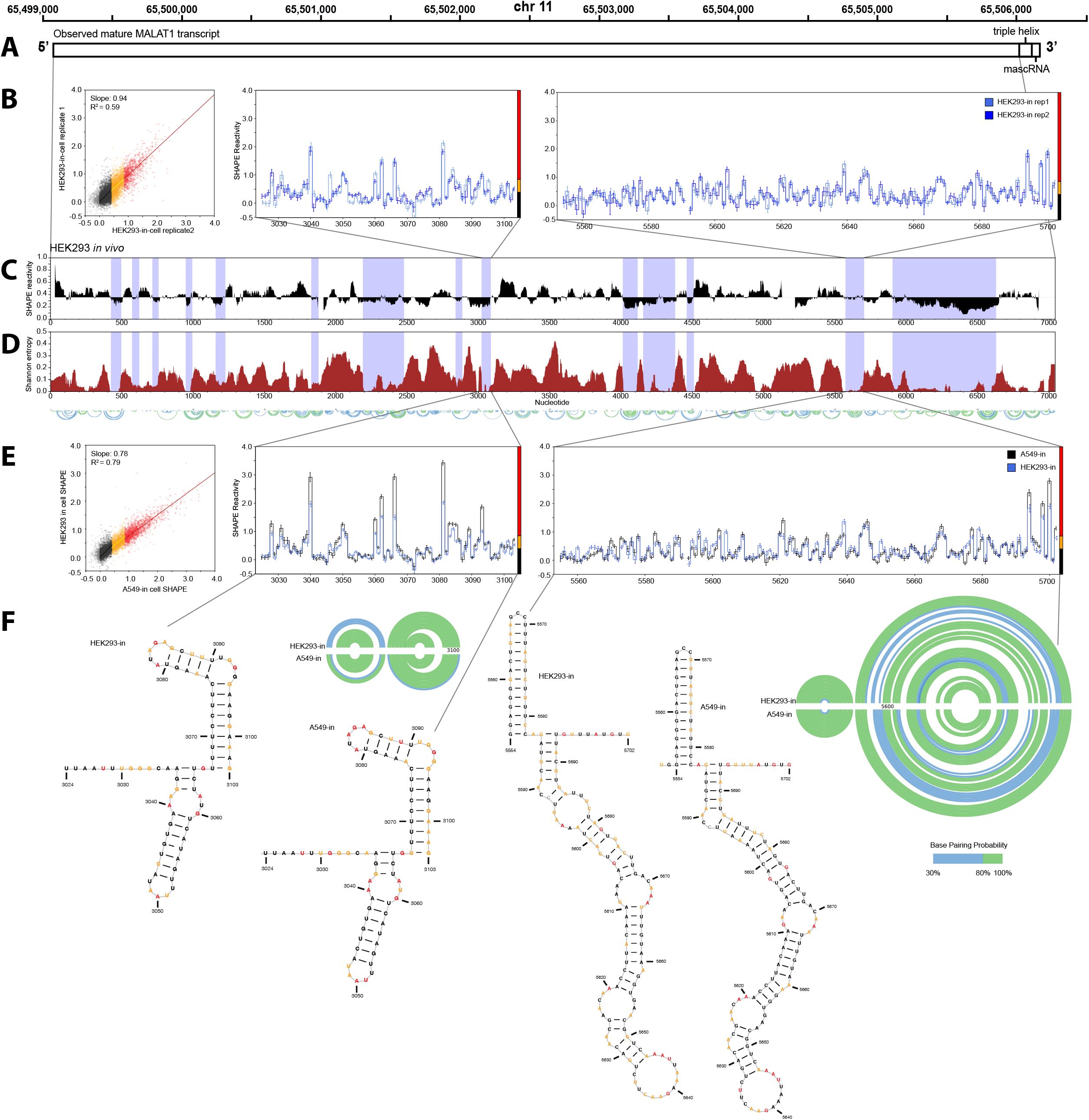
human MALAT1 RNA structure is nearly identical in cell in two different cell lines. A) Schematic of expressed MALAT1 transcript in human (GRCh38) genome coordinates observed in both A549 and Hek293 cells. B) in cell SHAPE-MaP data is highly reproducible across replicates in Hek293 cells. C) Median windowed (50 nt) SHAPE reactivity for Hek293 in cell data D) SHAPE derived Shannon Entropy identifies similar regions of high structure in Hek293 cell lines. E) Correlation analysis of SHAPE reactivities colored by low (black), medium (yellow) and high (red) reactivity show’s high correlation between Hek293 and A549 cells. Interestingly these are highly correlated, albeit with a slope of 0.78, suggesting the 5NIA can more easily access the RNA in A549 cells. This is also visible in comparison plots of reactivities where the overall pattern, which indicates structure, is very similar. F) When we model base pairing probabilities (arc diagrams) and corresponding minimum free-energy structures of low SHAPE, low Shannon entropy regions in the MALAT1 mRNA we obtain identical structures for both cell types. This suggests that the cellular environment is not significantly altering the structure of the MALAT1 mRNA in these two cell types and that the observed structures fold into similar conformations despite a different cellular environment.

### Structural differences of MALAT1 lncRNA in the Green Monkey Vero cell

Given the surprising quantitative consistency of our data across different experimental conditions, we now investigate the structure of MALAT1 in the African green monkey, *Chlorocebus sabaeus*, also known as the vervet using the Vero cell line (ATCC CCL 81, derived from kidney tissue of from adult African green monkey). The African green monkey and human share a common ancestor from approximately 25 million years ago (Kumar and Hedges 1998). As with Hek293 and A549 cells, we performed RNA-seq in Vero cells and identified the region of the transcript that is expressed (Figure 4A). As can be seen in Supplementary Figure 2, the Vero MALAT1 transcript is not spliced and has very similar primary structure to the human cell lines examined in this study, including a similar length. Since MALAT1 is unspliced in Vero cells as well, we were able to align Vero transcripts to the *Chlorocebus sabeus* genome (GenBank Accession GCA_000409795.2) to chromosome 1. We used genomic sequence that were aligned to human genomic coordinates for MALAT1 (Karolchik et al. 2008; Pollard et al. 2010). Currently, full length MALAT1 is not annotated in the vervet transcriptome, and, to our knowledge, we are the first to report the full-length vervet MALAT1 transcript. We were also able to observe the putative conserved triple helical region (Supplementary Figure 2), based both on sequence conservation and lower read depth (as we also observed in human). This is illustrated in Figure 4A, and we define the observed Vero transcript in Figure 4B.

**Figure 4:**
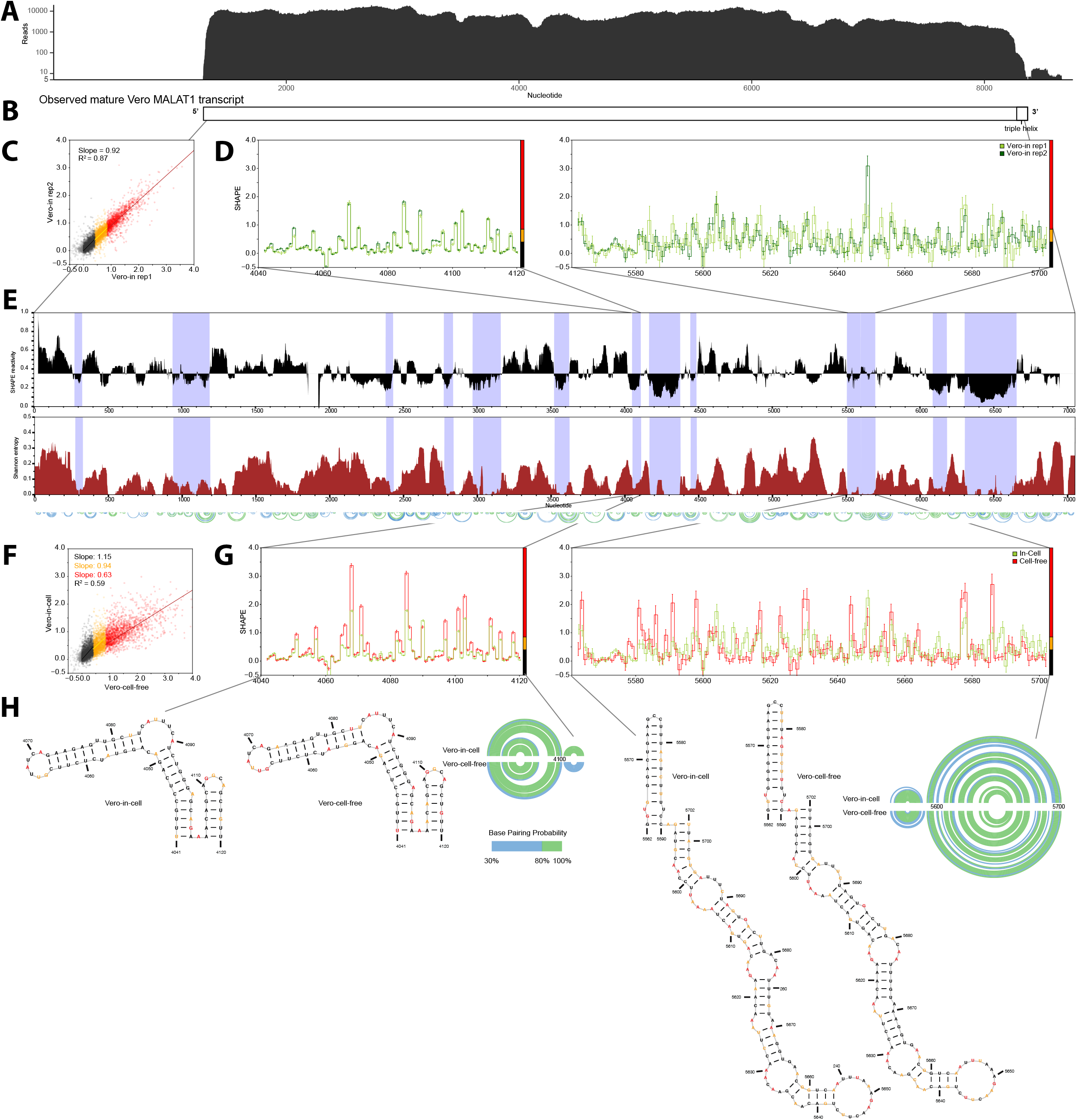
Structure of MALAT1 in green monkey Vero cells in vs. cell free reveals global changes in reactivity with minimum effect on highly structured regions of MALAT1. A) RNA-seq coverage of expressed MALAT1 transcript in Vero cells from which we determined the B) observed transcript isoform in these cells. C) in cell SHAPE-MaP data is highly reproducible across replicates as can also be seen in D) representative traces of SHAPE data for two regions in MALAT1. E) Median windowed (50 nt) SHAPE reactivity for in cell data combined with SHAPE derived Shannon Entropy. F) Correlation analysis of SHAPE reactivities colored by low (black), medium (yellow) and high (red) reactivity shows a clear decrease in slope for highly reactive nucleotides, despite an overall correlation comparable to in cell replicates. This is also apparent in G) representative regions of SHAPE data. H) When we model base pairing probabilities (arc diagrams) and corresponding minimum free-energy structures of low SHAPE, low Shannon entropy regions in the MALAT1 mRNA we obtain very similar structures both in and cell free. This is consistent with what we observed in human cell lines.

As with previous data sets reported in this manuscript, we observe very high replicate in cell correlation overall (Figure 4C) and for specific regions as well (Figure 4D). We used similar primers for the amplicons as those used in human cell lines (supplementary table 1) in these Vero cell experiments (supplementary table 2). From these we compute median SHAPE and Shannon entropy to identify low SHAPE, low Shannon entropy regions (Figure 4E). We collected cell free MALAT1 data in the Vero cell lines, and when we compared these with the in cell data (Figures 4F and 4G) we observed the similar decrease in slope in the high reactive (red) nucleotides, consistent with non-specific shielding of the cellular environment. The overall correlation in the data (R^2^=0.59) is still good, and the SHAPE patterns of in cell (green) and cell free (red) are very similar (Figure 4G). Similarly, when we model the low-shape, low Shannon entropy regions using these data, we obtain nearly identical base-pairing probabilities and minimum free energy structures (Figure 4H). Using these data, we will now investigate how substitutions from human to green monkey have affected the structure of these two MALAT1 lncRNAs.

### Structural comparisons of MALAT1 lncRNA in the Green Monkey and Human

To investigate the structural changes caused by substitutions in the MALAT1 transcript we aligned both SHAPE data sets to a consensus sequence (Figure 5A). In doing so we were able to identify 335 substitutions (differences between the human and green monkey transcripts), comprised of 140 transitions, 124 transversions, and 71 insertions and deletions. These substitutions are indicated using vertical white lines in Figure 5B. As is apparent from this analysis the substitutions are not evenly distributed but appear throughout the transcript. In Figure 5C we created a mountain plot where the y-axis is the number of substitutions in a 50-nucleotide window. Certain mountains peak at over 15 substitutions per 50 nucleotide window (e.g. near nucleotide 2000) with an effective divergence rate of 30% (*i*.*e*., 0.3 substitutions per site), while other regions (e.g. centered in nucleotide 4250) have divergence rate around 2%.

**Figure 5:**
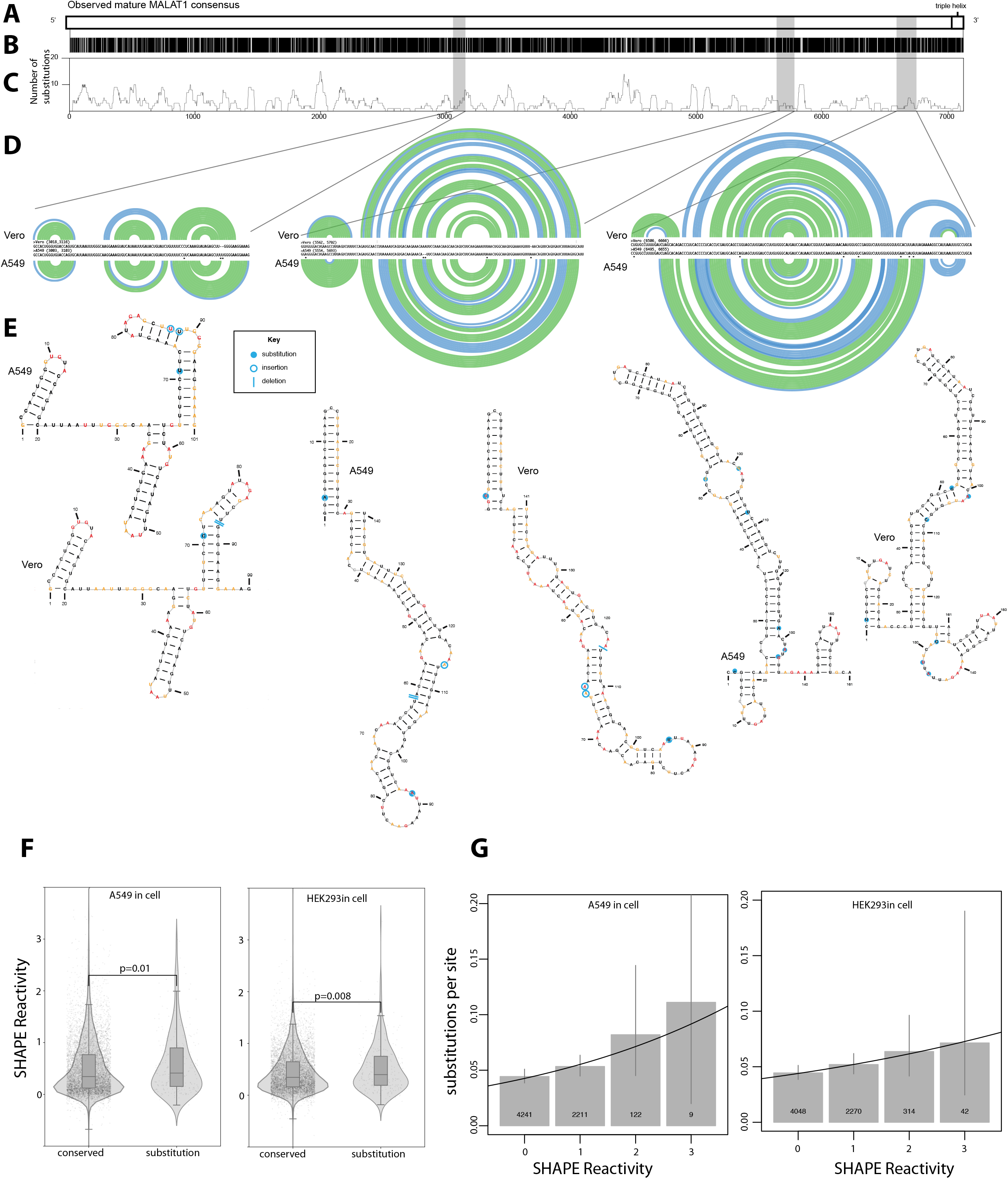
Comparative structural analysis of human and green monkey MALAT1 lncRNAs. A) Consensus MALAT1 transcript observed in all cell lines. B) Substitution analysis of human and green monkey MALAT1 lncRNAs, vertical white lines indicate sites of substitution, insertion or deletion. C) Mountain plots of substitutions using a 50-nucleotide window, where the y-axis is the number of substitutions in the window, illustrating regions of high substitution. D) Arc plot diagrams including alignments of Vero and A549 base-pairing probabilities for three representative low-SHAPE low Shannon entropy overlapping regions. Sequence alignment is shown, and sites of substitution are indicated with an *. These indicate a high level of structural conservation, which can be seen in E), the corresponding minimum free energy structures. Sites of substitution, insertion and deletion are indicated with blue labels on the structures. F) SHAPE reactivity analysis of sites with vs. without substitutions for the in-cell data sets reveal a higher median SHAPE for sites with substitutions residues, suggesting that conserved sites are more likely to be paired. G) The best-fit logistic regression of substitution presence/absence as a function of SHAPE reactivity (solid black lines) estimated that substitution rate increased 131% (Hek293) and 120% (A549) with every unit increase in SHAPE reactivity. Gray bars show the proportion of sites with substitutions (± 95% confidence intervals) for sites with SHAPE reactivities less than 0.5, 0.5 – 1.5, 1.5 – 2.5, and greater than 2.5. The number of sites within each bin is shown at the bottom of the bar.

SHAPE-directed structural modeling was optimized using reference RNAs with well-defined structures. These reference structures generally had low median SHAPE reactivity and low Shannon entropy (Deigan et al. 2009; Hajdin et al. 2013). As such, we can only compare SHAPE-directed structures for regions of low-SHAPE, low Shannon entropy that overlap between human and green monkey. In Figure 5D we show arc probability plots for three of these regions, which also have intermediate divergence rates. We used the in cell SHAPE data to model these structures to capture the most biologically relevant conformations, although these regions fold very similarly cell free.

What is immediately apparent from the base-pairing probability plots in Figure 5D is their high degree of similarity. Indeed, when we analyze the SHAPE-directed minimum free energy structures we also observe similar folds (Figure 5E). It is important that not all elements of the secondary structure are identical though, and the structures do differ in specific helices. Overall substitutions in unpaired regions appear to not alter the RNA structures, but those mapping to paired regions do. Thus, the question arises as to whether paired on unpaired nucleotides have a different rate of substitution.

We first used a t-test to compare the SHAPE reactivity of conserved positions (identical nucleotide in humans and green monkeys) versus substituted positions (different nucleotide in humans than in green monkeys) in the MALAT1 transcript. Consistent with the structure conservation, conserved positions exhibited significantly lower SHAPE reactivities than substituted positions (measured in Hek293: 0.050 difference in mean SHAPE reactivity, *t* = 1.92, df = 342, *p* = 0.028; measured in A549: 0.057 difference in mean SHAPE reactivity, *t* = 1.70, df = 348, *p* = 0.045; using one-sample t-tests), Figure 5F. Thus, conserved positions are more likely to be paired than substituted positions. Flipping the causative relationship around, we then used logistic regression to investigate the effect of SHAPE reactivity on the rate of nucleotide substitutions. The logistic regression analysis estimated a 131% (or 120%) increase in substitution rate with every increase in SHAPE reactivity of 1 unit as measured in the Hek293 (or A549) cell-lines (Figure 5G and Table 1). Thus, the least-paired positions (SHAPE ≥ 3), exhibit substitution rates that are at least 1.31^3^ = 2.25 (or 1.2^3^ = 1.73) times higher than the most-paired positions (SHAPE ≤ 0).

**Table 1:** Logistic regression estimates of the effect of SHAPE reactivity on substitution rate.

|  | **Estimate** | **Standard Error** | **Z value** | **Pr(>\|z\|)** | **Odds Ratio\*** |
| --- | --- | --- | --- | --- | --- |
| **A549 in cell reactivity** | 0.178 | 0.09 | 1.85 | 0.065 | $e^{0.178}=1.2$ |
| **Hek293 in cell reactivity** | 0.273 | 0.13 | 2.12 | 0.034 | $e^{0.273}=1.3$ |

\* The Odds Ratio is the proportional increase in the rate of substitutions per 1 unit increase in SHAPE reactivity.

## DISCUSSION

In this manuscript we investigated the structure of the lncRNA MALAT1 expecting to detect regions where either the cellular environment or evolutionary substitutions significantly altered the structure. It is likely lncRNA function is cell-type specific, since they are often expressed at low levels in a tissue specific manner (Cabili et al. 2011). In fact, RNA structure in general is impacted by the cellular environment (Rouskin et al. 2014; Tomezsko et al. 2020). This suggests that it is also important to understand the effect the environment has on the lncRNA and the different pressures that may be affecting how it folds in the cell. Previous studies have shown that it is difficult to identify even homologs of lncRNAs past a relatively short evolutionary distance (Necsulea and Kaessmann 2014; Necsulea et al. 2014; Hezroni et al. 2015), so we focused our attention on an exceptional lncRNA that is known to have function and is also known to be conserved in many different species. Albeit structural conservation is seen only at the 3’ end outside of mammals (Brown et al. 2014; Xiping et al. 2018). MALAT1 is an abundant nuclear RNA, with roles in modulating gene expression by binding to transcription factors and other RNAs (Wilusz et al. 2008; Brown et al. 2012; Wilusz et al. 2012). It also is known to bind to many additional proteins (Spiniello et al. 2018; Scherer et al. 2020).

We used SHAPE-MaP and a gene specific primer amplification strategy to examine full length MALAT1 structure in cells and cell free, in two cell types, and across two species (African green monkey and Human). First, we demonstrated high reproducibility between replicates in all cells studied. For each cell line, we observed that MALAT1 structure was very similar whether probed in cell or cell free. When we compared A549 cells to Hek293 cells, anticipating a remodeling of structure in the different in cell contexts, we saw that MALAT1 structure remained largely unchanged. This result is especially surprising given MALAT1’s conferral of metastatic potential in specific lung cancer cell lines, including A549 cells (Gutschner et al. 2013). Finally, we were curious if MALAT1 structure would be impacted by evolutionary substitutions that accrue between species. When we compared MALAT1 structure models derived from in cell probing of either a human cell line or a primate cell line, despite differences in sequence, we saw structure was still conserved, especially in common low-SHAPE, low-Shannon entropy regions.

Our metric for structural conservation is a very global one in this context. We chose to simply measure the correlation between our SHAPE data in different conditions. We chose to use this approach as opposed to compare SHAPE-directed structural models as these are notoriously sensitive to small differences in the data (Kladwang et al. 2011). This is especially true for large RNAs like MALAT1. When we focus on regions of low-SHAPE, low-Shannon entropy, the similarity between these high-confidence structural models continue to support our hypothesis. We considered computing windowed correlation coefficients to detect regions of high structural divergence. However, we found that this approach was very sensitive to small changes in the data and picking the correct window to use was arbitrary.

There are important caveats to this work. The fact that lncRNA MALAT1 has low-SHAPE, low Shannon entropy structures that are conserved does not necessarily mean that those structures are functional. They may simply appear conserved because there has not been sufficient evolutionary drift to disrupt them. Furthermore, in high SHAPE regions the structures likely form an ensemble. Nonetheless it is interesting that within experimental error, even in these “unstructured” regions there is strong correlation across cell types and species.

Of course, it is possible that SHAPE chemical probing is simply not sufficiently sensitive to capture subtle changes in structure. Finally, we focused on a relatively short evolutionary distance, anticipating that sequence differences would have a large impact on structure. Instead, we saw remarkable conservation. It will be important to examine the role of structure across longer evolutionary distances, and an obvious next step would be to probe the murine MALAT1 lncRNA. However, mouse MALAT1 sequence is significantly diverged from human, making direct alignments more complex. Thus, an intermediate evolutionary distance between monkey and mice could also be interesting (for example, marmoset and lemur).

## Materials and Methods

### Cell Culture

A549, Hek293, and Vero cells (ATCC CCL 185, CRL 1573, CCL 81, respectively) were cultured on tissue culture plates in DMEM supplemented with 10% Fetal Bovine Serum (FBS) and 0.5% penicillin and streptomycin at 37°C and 5% CO_2_. Experiments were performed before cells were completely confluent.

### In cell RNA SHAPE Treatment and RNA Extraction

Cells were grown in 10 cm plates and grown to 80-90% confluency. Cells were washed with PBS and replenished with 1350µL of media supplemented with 200mM bicine (pH 8.0) buffer (final concentration). 150µl 5-nitroisatoic anhydride (5NIA) at 250mM in dimethyl sulfoxide (DMSO) was added. Controls were treated with 150µL DMSO. Cells were then incubated at 37°C for 10 minutes. RNA was extracted by TRIzol purification and DNase digestion (ThermoFisher TRIzol, 5PRIME PhaseLock Heavy, Invitrogen Purelink RNA columns, ThermoFisher Purelink DNase Set) and quantified with NanoDrop™ spectrophotometer.

### Cell free RNA SHAPE treatment

Cell free experiments were performed on the same cell lines described above on natively transcribed RNA. RNA was Trizol extracted and quantified (as above). 10µg RNA from each experiment was incubated at 37°C for 10 min in folding buffer containing 100 mM Na-HEPES, pH 8.0, 100 mM NaCl, and 10 mM MgCl_2_. The RNA was incubated at 37°C for 5 minutes with 10% DMSO or with 250mM 5NIA in DMSO. Columns were used to purify the modified RNA (Invitrogen Purelink RNA columns).

### Reverse transcription and library preparation

For each sample of RNA, 3-10 µg was reverse transcribed using SHAPE-MaP reverse transcription with SuperScript II, random nonamers and low-fidelity conditions for all samples (Siegfried et al. 2014; Smola et al. 2015). (ThermoFisher Scientific SuperScript II, NEB random nonamers). The samples were purified with Ampure XP beads to isolate the cDNA and samples were eluted to 30µl. We designed species specific MALAT1 tiling primers for human and green monkey (Table S1) and performed multiplex PCRs following Qiagen Multiplex PCR protocol (Qiagen). cDNA was divided equally between primer sets. We performed secondary PCR to add TruSeq barcodes (NEB Q5 HotStart). Libraries were sequenced as paired end 2 × 250 read multiplex runs on a MiSeq instrument.

### SHAPE data analysis and RNA structure prediction

Two experimental repeats were collected for each cell type and condition. For the cell free data it included two Hek293 cell free repeats and a single A549 in cell experiment. SHAPE reactivities were derived using the ShapeMapper2 pipeline (Busan and Weeks 2018). A549 in cell repeats were combined; each reactivity for each nucleotide in A549 both in cell replicate experiments were combined and averaged to yield an averaged reactivity for each nucleotide. To compare data across experiments, each data set was averaged and combined (as A549 in cell above) and normalized to A549 in cell combined data. We normalized by finding the median SHAPE reactivity for each experiment and adjusting to match A549 in cell combined median by multiplying each value in that condition by the ratio of the A549 in cell shape median over the shape median for each specific condition. For structure diagrams, median shape, and Shannon entropy, the repeats for each condition were combined and averaged and normalized to A549 in cell data and are shown. For human cell free data, one replicate each of A549 cell free Hek293 cell free were combined and averaged and normalized as above. Arc plots, and graphs for median shape reactivity and Shannon entropy were obtained from SuperFold, which also incorporates chemical probing data (Smola et al. 2015). SuperFold was also used to find regions of significant structure in each of the conditions and was used to generate minimum free energy secondary structure models. SuperFold uses RNAstructure to calculate MFE, again, with chemical probing data incorporated (Deigan et al. 2009; Wilkinson et al. 2009). Secondary structure plots were created with Varna (Darty et al. 2009). Reactivity plots and scatter plots were created in Python using the PyPlot package.

### MALAT1 Conservation analysis and alignments and genome reference

We downloaded PhyloP-scored alignments of 30 mammals from UCSC Genome Browser http://hgdownload.cse.ucsc.edu/goldenPath/hg38/phyloP30way/ (Karolchik et al. 2008; Pollard et al. 2010) and show the PhyloP scores of genome coordinates coordinating with MALAT1. Since the full length MALAT1 transcript is not annotated, we used genomic African green monkey sequence (GenBank Accession GCA_000409795.2) that was aligned to human MALAT1 from the PhlyoP alignments above to align our RNAseq reads from vervet MALAT1 pull-down (Supplemental Materials and Methods). We were able to align our RNAseq reads to genomic African green monkey and report MALAT1 expression in Vero cells and found Vero cells expressed a transcript with similar primary structure and length to human Hek293 and A549 cells. ClustalOmega was used to align our human and vervet MALAT1 sequences to find a consensus sequence (Sievers and Higgins 2014; Sievers and Higgins 2021). The sequences were then compared, and at each point where they differed, a substitution was logged. Substitution types were classified into 4 different groups: transition or transversion if a nucleotide was present in both sequences, and deletion or insertion if only one sequence had a nucleotide at that position according to the alignment (insertion and deletion were categorized relative to the human sequence). At each position in the sequence, SHAPE data was logged and categorized according to whether a substitution was present at that position, and then further split by type. All data used during this process comes from the combined in cell experiments.

For each cell type (Hek293, A549, and Vero), the resulting datasets for conserved (human and green monkey sequences are identical) and diverged (human and green monkey sequences differ) positions were compared. Violin, strip, and box plots were generated in the Seaborn package for Python. t-tests were performed using the *t*.*test* function from the *stats* package in R version 2.2.0. Logistic regressions were performed using the *glm* function from the *stats* package in R, using a binomial error distribution and a logit link function. IGV viewer was used to as a reference to visualize MALAT1 coordinates in the human genome hg38 (Robinson et al. 2010).

### Data Access

Raw sequence reads are available at the sequence read archive (SRA accession number SUB11086382), SHAPE reactivities are provided in the supplemental material as a SNRNASM file, and a Github page https://github.com/LaederachLab/malat1 was created to facilitate access to the code used to analyze the data.

## Acknowledgments

This work was supported by U.S. National Institutes of Health grants NHLBI R01 HL111527 and NIGMS R35 GM140844 to A.L. A.M.E. was supported by the National Science Foundation Graduate Research fellowship under grant number 5105482. C.W. is supported U.S. National Institutes of Health grant K22CA262349.

## Notes

### Competing Interest Statement

The authors have declared no competing interest.

